# Cathepsin Z is a conserved susceptibility factor underlying tuberculosis severity

**DOI:** 10.1101/2025.04.01.644622

**Authors:** Rachel K. Meade, Oyindamola O. Adefisayo, Marco T. P. Gontijo, Summer J. Harris, Charlie J. Pyle, Kaley M. Wilburn, Alwyn M. V. Ecker, Erika J. Hughes, Paloma D. Garcia, Joshua Ivie, Michael L. McHenry, Penelope H. Benchek, Harriet Mayanja-Kizza, Jadee L. Neff, Dennis C. Ko, Jason E. Stout, Catherine M. Stein, Thomas R. Hawn, David M. Tobin, Clare M. Smith

## Abstract

Tuberculosis (TB) outcomes vary widely, from asymptomatic infection to mortality, yet most animal models do not recapitulate human phenotypic and genotypic variation. The genetically diverse Collaborative Cross mouse panel models distinct facets of TB disease that occur in humans and allows identification of genomic loci underlying clinical outcomes. We previously mapped a TB susceptibility locus on mouse chromosome 2. Here, we identify cathepsin Z (*Ctsz*) as a lead candidate underlying this TB susceptibility and show that *Ctsz* ablation leads to increased bacterial burden, CXCL1 overproduction, and decreased survival in mice. *Ctsz* disturbance within murine macrophages enhances production of CXCL1, a known biomarker of TB severity. From a Ugandan household contact study, we identify significant associations between *CTSZ* variants and TB disease severity. Finally, we examine patient-derived TB granulomas and report CTSZ localization within granuloma-associated macrophages, placing human CTSZ at the host-pathogen interface. These findings implicate a conserved CTSZ-CXCL1 axis in humans and genetically diverse mice that mediates TB disease severity.

## Introduction

*Mycobacterium tuberculosis* (*Mtb*), the causative agent of tuberculosis (TB), is a prolific obligate pathogen that has threatened human health for millennia [1]. Through centuries of co-evolution, human hosts have developed a plethora of immunological mechanisms in response to *Mtb* infection [2]. Such host-bacterial interactions give rise to a spectrum of disease states, ranging from subclinical infection to fulminant disease [3]. The disease severity experienced by an individual is intricately connected to their genetic background. For example, monozygotic twins are at a demonstrably higher risk for TB concordance than dizygotic twins, highlighting shared genetic identity as a contributor to TB disease outcomes [4–6]. Human genome-wide association studies (GWAS) conducted in impacted geographic regions have also identified polymorphisms that modulate host TB immunity [7–13], indicating numerous immunological pathways involved in *Mtb* susceptibility.

One such gene is cathepsin Z (*CTSZ*), which has been associated with TB susceptibility in independent human studies conducted across Africa. *CTSZ* encodes a lysosomal cysteine protease with a known structure and several reported cellular functions [14–22]. The link between single nucleotide polymorphisms (SNPs) in *CTSZ* and human TB susceptibility was first established by sibling pair analysis in South African and Malawian populations and independent case-control studies in West Africa [23]. These findings were further validated in a South African case-control study [24] and in a Ugandan GWAS [25] and subsequent household contact study [26]. *CTSZ* is primarily expressed by monocytes and macrophages [27–30], and participates in central immune functions, including dendritic cell maturation [31] and lymphocyte propagation and migration [32,33]. Although *in vitro* work has been undertaken to study the role of CTSZ in macrophage-driven protection against mycobacteria [34,35], *CTSZ*-linked TB susceptibility has not been explored *in vivo*. The functional role of CTSZ during *Mtb* infection remains unknown, despite growing genetic evidence of its association with TB disease outcomes.

Studying the mechanisms that underlie *CTSZ*-linked susceptibility in humans is complex [36]. Humans are genetically outbred, and the low- and middle-income countries that harbor 80% of the global TB burden face challenges in and outside of the healthcare sector that complicate diagnosis, research, and treatment [37]. The connection between TB severity and host background is not uniquely human. Classic studies measuring survival in inbred mice infected with *Mtb* illustrate the heritability of TB susceptibility [38,39], establishing mice as tractable models that demonstrate the vital impact of host genetic background on *Mtb* pathogenesis. However, because inbred mice are nearly genetically identical within strain [40], studies leveraging standard inbred strains omit the contributions of natural host genetic diversity to TB pathogenesis. Recombinant inbred panels like the biparental BXD [41–44], and octoparental Collaborative Cross (CC) [45–47] systematically model host genetic variation, allowing insight into a spectrum of immune profiles without compromising the reproducibility of inbred strains [48]. We previously reported *Mtb* infection screens of BXD [49] and CC [50] recombinant inbred strains, leveraging these diverse mammalian panels to expand the range of known TB disease complexes and host-pathogen interactions modeled by mice. Using a quantitative trait locus (QTL) mapping approach across a cohort of 52 CC genotypes, we identified a QTL on chromosome 2 (174.29-178.25Mb) significantly associated with *Mtb* burden. Genetic inheritance from NOD/ShiLtJ (NOD), a CC panel founder, at the *Tuberculosis ImmunoPhenotype 5* (*Tip5*) QTL predicted elevated bacterial burden. CC strains that inherited the susceptible *Tip5* variant (*Tip5^S^*) from NOD succumbed to severe TB prior to the study endpoint. We therefore sought to determine whether genes found within the *Tip5* interval contribute to *Mtb* susceptibility in *Tip5^S^* CC strains.

Here, we show that CC strains harboring the *Tip5^S^* locus produce lower levels of CTSZ protein while exhibiting higher bacterial burden than B6 mice following aerosol infection, validating *Tip5* as a susceptibility locus from the large-scale CC cohort screen. We report the first *in vivo Mtb* infections of mice lacking *Ctsz* (*Ctsz^−/−^*). We find that *Ctsz* ablation on a B6 background results in increased *Mtb* burden and an increased risk of mortality following infection. Moreover, *Ctsz*^−/−^ mice overproduce CXCL1, a biomarker of active TB [51], at both acute and chronic timepoints. In *Ctsz*^−/−^ bone-marrow derived macrophages (BMDMs), we find that CXCL1 is rapidly induced following mycobacterial infection. Leveraging published transcriptional data from genetically diverse mice, humans, macaques, and zebrafish, we find cathepsin Z expression is highest in macrophages following infection. We combine these findings with recent data from a Ugandan patient cohort, highlighting 5 variants in *CTSZ* as correlates of TB severity. Finally, we identify the presence of CTSZ in CD68^+^ macrophages within patient-derived pulmonary granulomas, revealing that CTSZ is produced at the host-pathogen interface in human lungs. Collectively, this work establishes genetic variation in cathepsin Z as a determinant of TB disease outcomes and places human CTSZ in a vital position within the pulmonary microenvironment to impact TB outcomes.

## Results

### Comparative transcriptional analysis to prioritize candidate genes within the

Tip5 *locus* We previously reported the *Tip5* QTL (Chr2, 174.29-178.25Mb) as a TB susceptibility locus across the genetically diverse CC panel [50]. To identify gene candidates within *Tip5*, we leveraged published transcriptomic data from *Mtb*-infected mammalian lungs [52,53] (**Figure 1A**). Within the *Tip5* interval, cathepsin Z (*Ctsz*; also, cathepsin X or P) and zinc finger protein 831 (*Zfp831*) were significantly induced in the lungs of genetically heterogenous Diversity Outbred (DO) mice exhibiting progressive TB, characterized by elevated pulmonary *Mtb* burden and inflammation [52]. In rhesus macaques, animals with progressive TB disease produced significantly more *CTSZ* and *ZNF831* transcript in their lungs [52]. In the blood of patients with active TB, *CTSZ* transcription was significantly elevated while *ZNF831* transcription was significantly repressed [52,54]. In an additional lung transcriptomic study in inbred mice leveraging distinct *Mtb* strains and infectious doses, only 5 gene transcripts within *Tip5*, including *Ctsz*, were differentially regulated across all strains and doses [53]. Currently, there is no established association between human *ZNF831* SNPs and TB outcomes. Conversely, mutations in human *CTSZ* were previously associated with poorer TB outcomes [23,24,26]. From this analysis, *Ctsz* was identified as a lead candidate for further interrogation as a potential genetic cause of *Tip5*-linked TB susceptibility.

**Figure 1:**
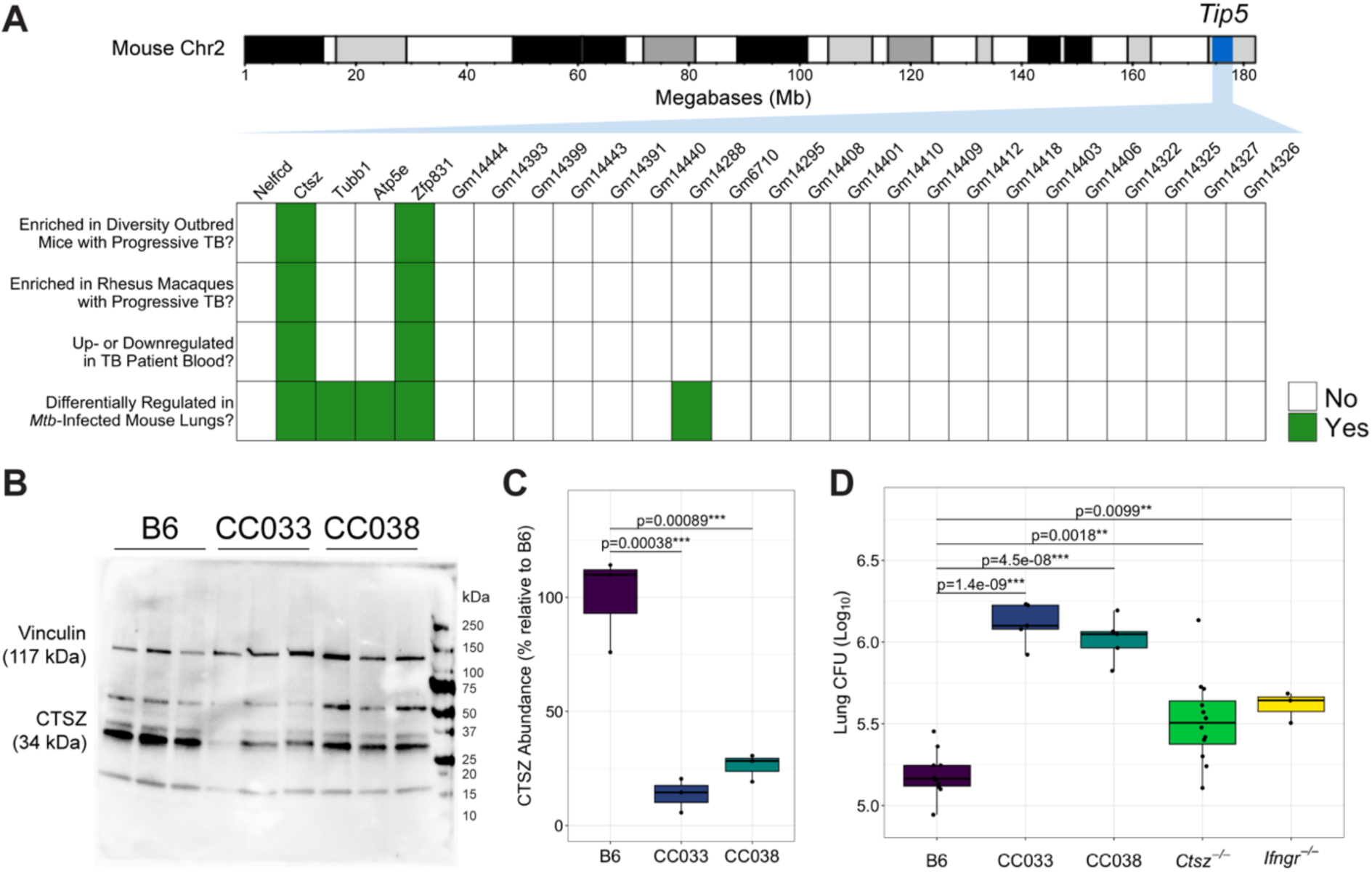
Identification and validation of *Ctsz* as the lead candidate gene underlying *Tip5*. **(A)** Heatmap representation of the per-gene outcome of four distinct criteria for genes within the *Tip5* QTL (95% CI: 174.29-178.25Mb): *i)* whether the gene transcript is significantly enriched in the lungs of genetically heterogeneous Diversity Outbred (DO) mice experiencing elevated burden and inflammation after *Mtb* infection [52], *ii)* whether the gene transcript is significantly enriched in the lungs of rhesus macaques exhibiting clinical symptoms of severe TB disease [52], *iii)* whether the gene is significantly up- or downregulated in the blood of individuals with active TB [54], and *iv)* whether the gene is differentially expressed in inbred mouse lungs across variable host genotypes, *Mtb* strains, and infectious doses [53]. To be included in the heatmap, genes were required to encode proteins and to contain a known SNP from the NOD inbred line [55]. Mouse chromosome 2 image generated in the R package karyoploteR. (**B**) CTSZ protein was measured from the lung homogenate of uninfected B6 and the *Tip5^S^>* CC strains CC033 and CC038 (n=3 mice per genotype). Each lane is a separate biological replicate. Vinculin served as the loading control. (**C**) Relative abundance of the CTSZ protein between B6 and the *Tip5^S^>* CC strains, quantified from Figure 1B by normalizing CTSZ levels for each biological replicate to its respective vinculin level. Values plotted as a percentage of the mean CTSZ to vinculin band intensity ratio relative to the average ratio for B6 mice. Hypothesis testing was performed by one-way ANOVA and Dunnett’s *post hoc* test on individual ratios between CTSZ and vinculin band intensities by genotype. (**D**) Bacterial burden measured from lung homogenate 4 weeks after aerosol infection with *Mtb* H37Rv (n=3-12 mice per strain; all males except B6 and *Ctsz^−/−^* groups, which included both sexes in equal proportion). Hypothesis testing was performed by one-way ANOVA and Dunnett’s *post hoc* test on log_10_-transformed values.

### The susceptible NOD variant of Tip5 and ablation of Ctsz both impart TB susceptibility

To evaluate *Ctsz* as a causal factor underlying *Tip5*-linked susceptibility, we measured CTSZ protein from the lungs of uninfected CC strains harboring the susceptible NOD *Tip5* variant (CC033, CC038). Compared to *Mtb*-resistant B6, the lungs of both CC033 and CC038 exhibited significantly lower baseline levels of CTSZ protein (**Figure 1B** & **C**). Collectively, these data suggest that the NOD *Tip5* haplotype contains a hypomorphic variant of *Ctsz*, resulting in reduced production of CTSZ protein in *Tip5^S^>* CC strains.

Considering the *Tip5* QTL was first identified in a large-scale *in vivo* screen, we next assessed whether *Tip5^S^>*CC strains and *Ctsz* null mice (*Ctsz*^−/−^) (**Figure S1A**-**B**) are susceptible to aerosol infection, the natural route of *Mtb* infection. A cohort including B6, CC033, CC038, *Ctsz^−/−^*, and highly susceptible interferon gamma receptor null mice (*Ifngr^−/−^*) [56] was infected via aerosol route with *Mtb* H37Rv. The experiment terminated at 4 weeks post-infection, after the onset of adaptive immunity [57] and matching the initial CC screen endpoint [50]. Relative to B6, all infected strains exhibited significantly higher pulmonary *Mtb* burden (**Figure 1D**). The CC strains exhibited tenfold greater lung CFU than B6, surpassing the canonically susceptible *Ifngr^−/−^* mice. *Ctsz^−/−^* mice exhibited a twofold increase in lung burden relative to B6. No significant differences were identified in disseminated spleen burden at this timepoint (**Figure S1C**). We conclude that *Tip5^S^>* CC strains and *Ctsz^−/−^* mice exhibit reduced pulmonary bacterial control at 4 weeks post-infection.

Ctsz *mediates lung CXCL1 levels early during* Mtb *infection*

To characterize the impact of *Ctsz* on disease progression, we infected B6 and *Ctsz^−/−^* mice via aerosol, sacrificing cohorts of mice at 2, 3, 4, and 8 weeks post-infection to capture innate and adaptive immune responses. *Ctsz^−/−^*mice exhibited higher lung burden at 2 weeks (4.09 log_10_ CFU vs. 3.41 in B6; p<0.05) and 4 weeks (5.17 log_10_ CFU vs. 4.09 in B6; p<0.05) post-infection (**Figure 2A**). Similarly, at 3 weeks post-infection, *Ctsz^−/−^* mice exhibited trends toward elevated spleen burden (2.68 log_10_ CFU vs. 2.17 in B6; p=0.058), suggesting earlier dissemination and weaker bacterial containment in the lungs of *Ctsz^−/−^* mice (**Figure 2B**). However, by 4 weeks post-infection, spleen burden was indifferentiable between *Ctsz^−/−^*and B6.

**Figure 2:**
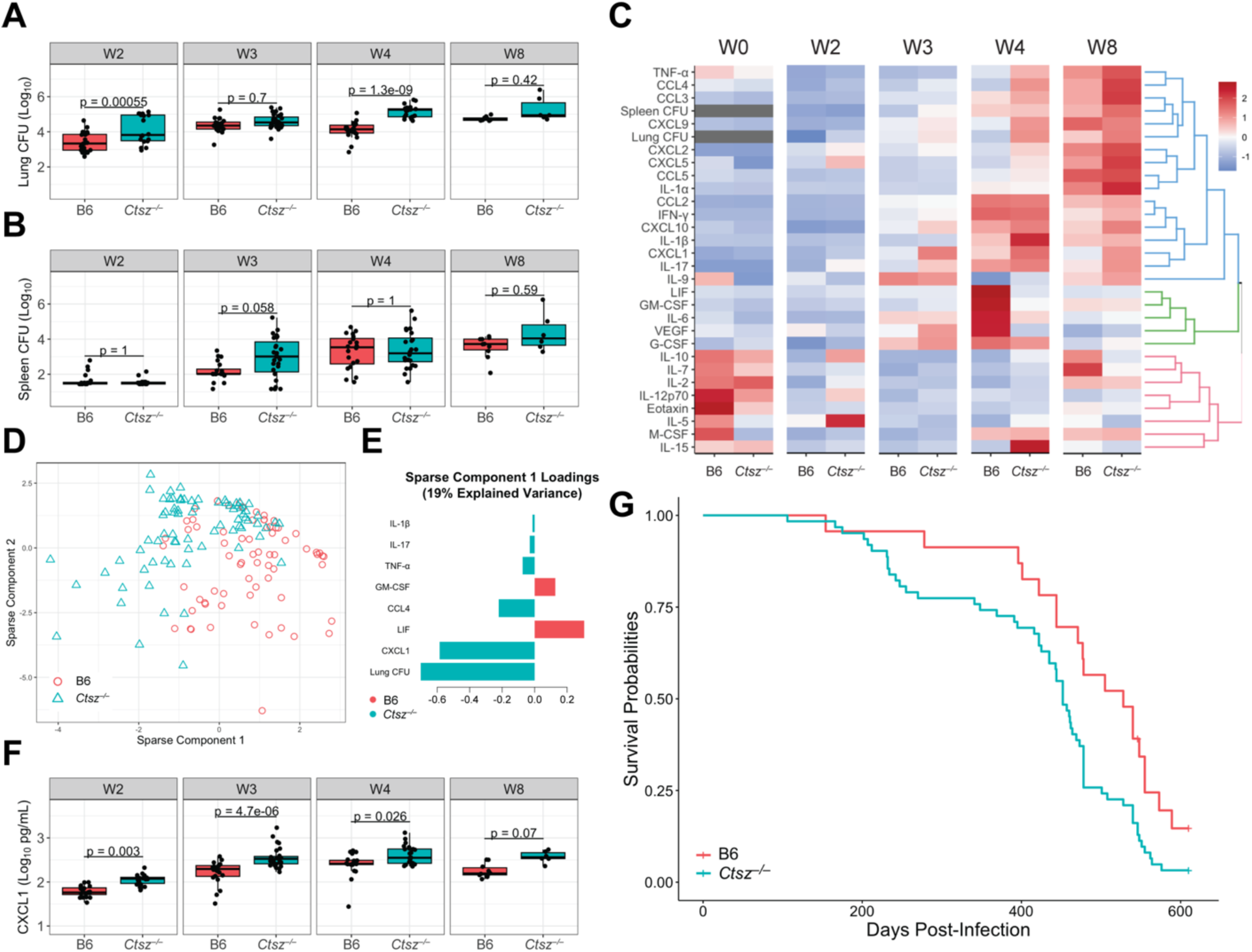
*Ctsz^−/−^*mice have higher lung burden and earlier spleen dissemination during acute infection followed by greater chronic inflammation and mortality risk. *Mtb* burden measured from (**A**) lung and (**B**) spleen homogenate by dilution plating. (**C**) Heatmap depicting scaled and centered phenotypes, hierarchically clustered and separated into 3 k clusters. (**D**) Individual mice plotted against the first two sPLS-DA components, which explained the greatest variance in the data after optimization. (**E**) Phenotype loadings contributing to component 1. Component 2 loadings shown in **Figure S2A**. (**F**) CXCL1 levels measured from lung homogenate by multiplex ELISA. For panels **A**, **B**, and **F**, hypothesis testing was performed by two-way ANOVA and Tukey’s *post hoc* test on log_10_-transformed values. For panels **A**-**F**, mice were sacrificed at 2, 3, 4, and 8 weeks following aerosolized *Mtb* H37Rv infection. Data are from two independent experiments with n=6-14 mice per genotype, representative of both sexes, at each timepoint. In panel **C**, age-matched, uninfected mice (n=3-4 per genotype and sex) were assayed for comparison. (**G**) Kaplan-Meier survival estimates of aerosol-infected B6 (n=23) and *Ctsz^−/−^* mice (n=62) across two independent experiments. Equal proportions of both sexes were included. Mice that were not moribund at time of sacrifice were censored for analysis.

To profile the impact of *Ctsz* disturbance on the lung inflammatory response throughout the course of infection, we compared cytokine signatures of *Ctsz^−/−^* with B6 at assayed timepoints. At 4 weeks post-infection, *Ctsz^−/−^* mice exhibited higher concentrations of T_H_1-associated cytokines, like TNF-α (p=0.019) and IL-1β (p=0.016), and lower levels of GM-CSF (p=3.8e-06), IL-6 (p=5.9e-04), LIF (p=6.6e-07), and VEGF (p=6.6e-07) compared to B6 (**Figure 2C**).

To identify the inflammatory signature of *Ctsz^−/−^* mice in an unsupervised manner, we performed sparse partial least squares discriminant analysis (sPLS-DA) across measured phenotypes (**Figure 2D**). Higher lung burden and CXCL1 levels in *Ctsz^−/−^* mice were the strongest features underlying sparse component 1 (**Figure 2E**). Although component 1 explains 19% of variance in the data compared to 23% variance explained by component 2 (**Figure S2A**), component 1 better captures the variance attributable to genotype. CXCL1 has previously been identified as a biomarker of active TB disease in genetically diverse mice [51] and in humans [58]. From 2 to 4 weeks post-infection, *Ctsz^−/−^* mice exhibited significantly higher lung CXCL1 levels (**Figure 2F**), suggesting that *Ctsz* ablation increases disease severity. However, by 8 weeks post-infection, although mean CXCL1 levels in *Ctsz^−/−^*lungs were elevated, the difference was no longer significant. Enhanced production of CXCL1 was consistent throughout infection, suggesting that this effect may occur independent of differences in *Mtb* burden.

To explore the possibility that elevated CXCL1 levels may occur independent of infection in *Ctsz^−/−^* mice, we sacrificed uninfected mice of both sexes. From lung homogenate, we found elevated levels of CXCL1 in *Ctsz^−/−^* compared to B6 (**Figure 2C**; p=0.007), suggesting that the connection between *Ctsz* and CXCL1 extends beyond the context of infection. Notably, the total CXCL1 levels in uninfected mice were comparable to levels measured at 2 weeks post-infection.

To determine whether *Ctsz* ablation alone is sufficient to confer susceptibility to aerosolized *Mtb* H37Rv, we conducted two longitudinal challenges of B6 and *Ctsz^−/−^* mice in which mice were sacrificed when IACUC-approved humane endpoints were reached. *Ctsz* ablation was associated with a significant reduction in survival time (p=0.008; **Figure 2G**), which was driven by male mice (**Figure S2B** & **C**). Thus, disease progression in a host lacking *Ctsz* is characterized by increased lung *Mtb* burden, lung CXCL1 levels, and overall mortality risk.

### Disturbance of Ctsz enhances CXCL1 induction in macrophages

To explore the expression of cathepsin Z across species and mycobacterial infection models, we analyzed two previously published single-cell RNA sequencing (scRNA-Seq) datasets. In zebrafish infected with *Mycobacterium marinum (Mm)*, *ctsz* was most highly expressed in inflammatory macrophages (cluster 9) after 14 days of infection (**Figure 3A** & **B**) [59]. *CTSZ* in cynomolgus macaques was most highly expressed in macrophages 4 weeks after *Mtb* infection compared to other assayed cell types (**Figure 3C** & **D**) [60]. These results agree with literature establishing the presence of CTSZ in monocytes and macrophages [27–30] and further highlight that cathepsin Z expression in these cell types following mycobacterial infection is conserved across diverse host species.

**Figure 3:**
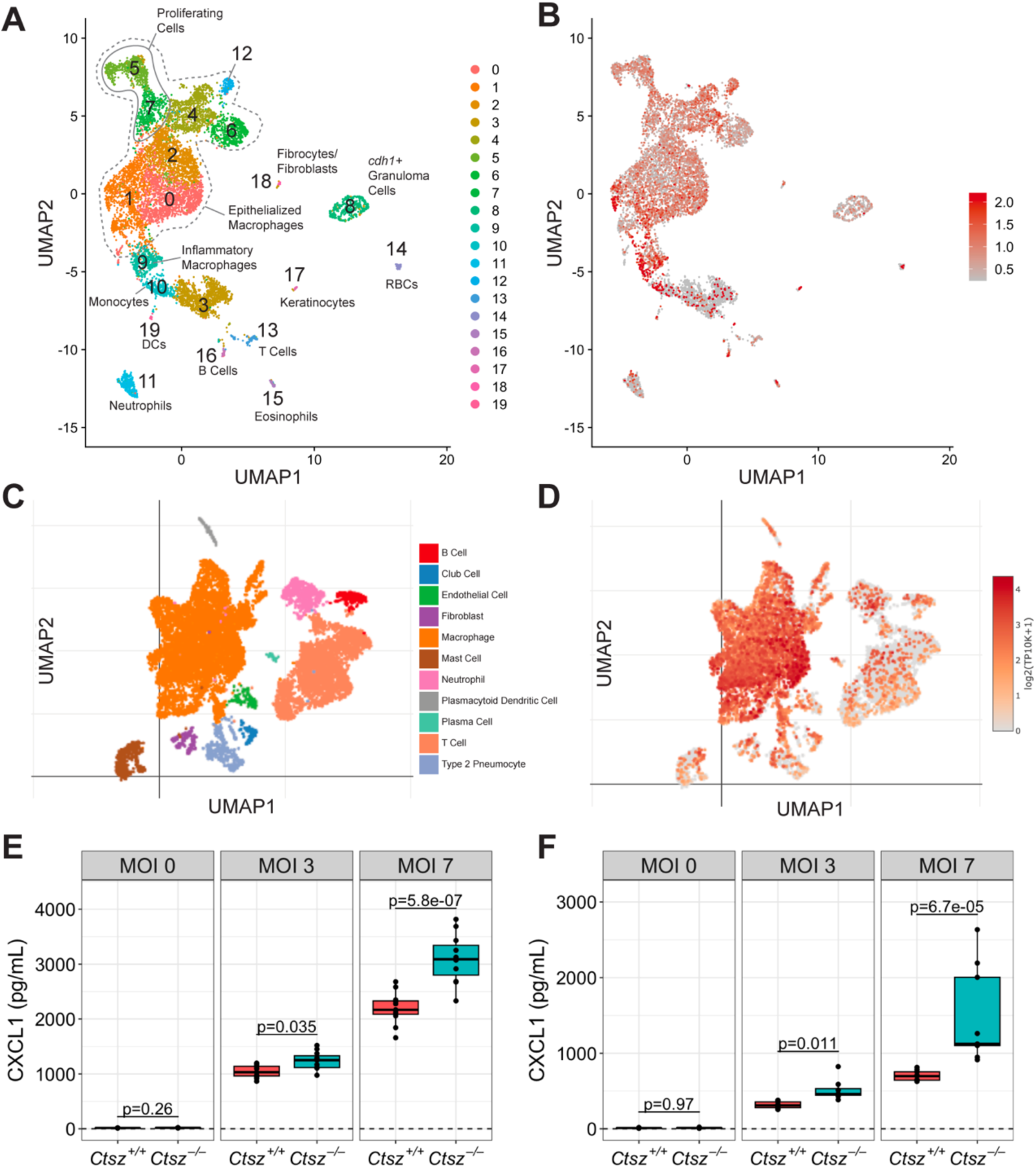
Cathepsin Z is highly expressed in macrophages across species following mycobacterial infection and mediates levels of CXCL1 in murine macrophages. (**A**) UMAP showing scRNA-Seq results from zebrafish granulomas infected with *M. marinum* (*Mm*), first published by Cronan, Hughes et al., 2021 [59]. Cells are colored by cluster and assigned in an unsupervised approach from transcriptional signatures. Clusters were annotated by the authors. (**B**) ScRNA-Seq data colored by relative expression of zebrafish *ctsz* in *Mm*-infected granulomas, with highest expression levels observed in cluster 9. (**C**) UMAP showing scRNA-Seq of granulomas extracted from *Mtb*-infected cynomolgus macaques at 4 weeks post-infection, published by Gideon et al., 2022 [60]. Data accessed at the Broad Institute Single Cell online repository on October 3, 2023 (https://singlecell.broadinstitute.org/single_cell/study/SCP1749/). (**D**) ScRNA-Seq data colored by relative expression of *CTSZ* in *Mtb*-infected macaque granulomas. CXCL1 levels measured in triplicate at 24 hours post-infection from (**E**) BCG-infected and (**F**) *Mtb*-infected BMDMs by ELISA. For panels **E** and **F**, BMDMs were differentiated from independent pairs of *Ctsz^+/+^>* and *Ctsz^−/−^* sibling males for each infection (N=3 infections per pathogen). Dashed threshold denotes the limit of detection for the ELISA. Statistical significance was determined by two-way ANOVA and Tukey’s *post hoc* test on batch-corrected, log_10_-transformed values.

As cathepsin Z is consistently expressed in macrophages across several species following mycobacterial infection, we sought to characterize the impact of *Ctsz* ablation on the initial macrophage response to mycobacterial exposure. To test if macrophages contribute to the increased production of CXCL1 during infection in *Ctsz^−/−^* mice, we generated BMDMs from *Ctsz^+/+^>* and *Ctsz^−/−^* sibling pairs. When infected with either non-pathogenic *Mycobacterium bovis* (Bacillus Calmette-Guérin; BCG) (**Figure 3E**) or *Mtb* (**Figure 3F**), *Ctsz^−/−^* macrophages produced greater amounts of CXCL1 than *Ctsz^+/+^>* by 24 hours post-infection. In both infection models, this effect scaled with increasing multiplicity of infection (MOI). Thus, the elevated CXCL1 we observed in *Ctsz^−/−^* lungs may be driven by macrophages, especially during the early stages of infection, and appears to be independent of mycobacterial pathogenicity. These results from *Mtb*-infected *Ctsz^−/−^*mice and BMDMs suggest an interaction between CTSZ and CXCL1 following bacterial exposure.

### Variants in human CTSZ are associated with TB severity

To investigate the impact of natural *CTSZ* variation on human TB disease outcomes, we examined whether human *CTSZ* variants are associated with TB disease severity in a household contact study in Uganda (n=328 across two independent cohorts) [61]. Of 81 observed *CTSZ* SNPs, 20 SNPs were associated with differences in Bandim TBScore, a TB severity index (**Table S1**; unadjusted p<0.05, linear model with sex, HIV status, and genotypic principal components 1 and 2 as covariates) [62]. After performing a Bonferroni adjustment for multiple comparisons, 4 SNPs and 1 INDEL maintained associations with TB disease severity (**Table 1**). These variants are in strong linkage disequilibrium (LD) with one another (R^2^>>0.8), suggesting that they represent a single haplotype block (**Figure 4A**). For the most significant SNP (rs113592645), the minor T allele is associated with decreased TB disease severity (**Figure 4B**, results for other haplotype SNPs included in **Figure S3A**-**C**). To investigate the potential impact of the TB severity SNPs on *CTSZ* expression, we used published RNA-Seq data [63] to compare *CTSZ* transcript levels across *Mtb*- and mock-infected monocytes between genotypes at each *CTSZ* SNP. In the cohort of human-derived monocytes, *CTSZ* was highly expressed at baseline and was downregulated following *Mtb* infection (**Figure 4C**). Conversely, the rs113592645 minor T allele was associated with increased *CTSZ* expression following *Mtb* infection (p=0.0395; **Figure 4D**; other haplotype SNP results in **Figure S3D**-**F**). This effect was not observed following mock infection conditions (p=0.108; **Figure 4D**). Together, these data suggest that *CTSZ* variants are associated with both TB disease severity and divergent transcription of *CTSZ* following *Mtb* infection.

**Figure 4:**
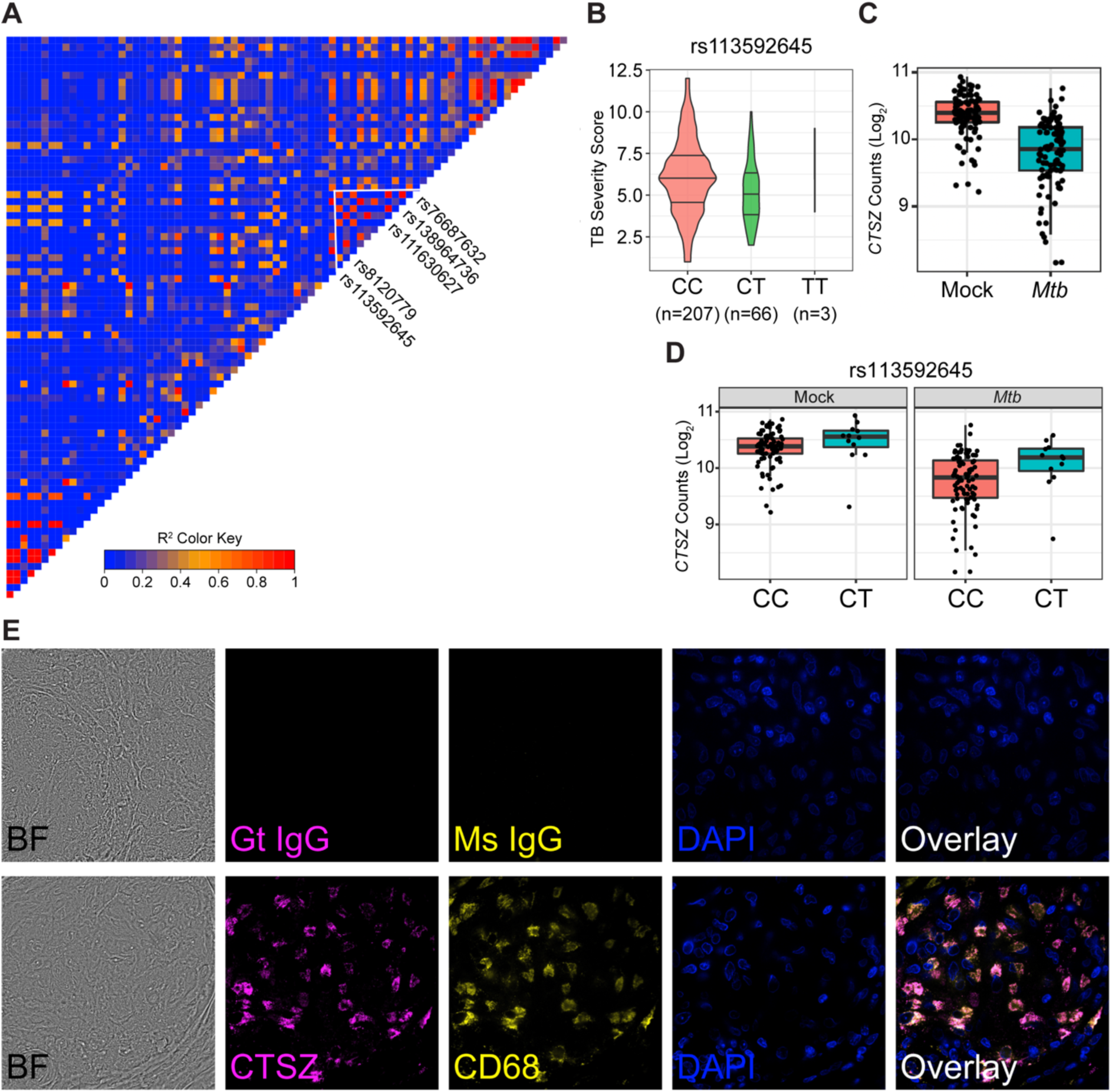
Human *CTSZ* variants are associated with TB severity, and CTSZ protein is produced within the human pulmonary granuloma. (**A**) LD plot of human *CTSZ*, highlighting a haplotype block of 4 identified SNPs and 4 SNPs and 1 INDEL associated with TB disease severity. (**B**) Comparison of TB severity, measured using Bandim TBScore, by genotype for the lead TB severity SNP, rs113592645. TB severity score by genotype for remaining SNPs can be found in **Figure S3A**-**C**. For panels **C** and **D**, *CTSZ* expression was profiled by RNA-Seq in monocytes from 100 Ugandan individuals. Human-derived monocytes were subjected to 6-hour *Mtb* and mock infection conditions. (**C**) Counts of *CTSZ* transcript (log_2_ counts per million) collected following mock and *Mtb* infection. (**D**) Counts of *CTSZ* transcript (log_2_ counts per million) according to genotype for the lead TB severity SNP, rs113592645, following mock and *Mtb* infection. Measurements for homozygous minor allele (TT) were excluded due to low sample size. Counts of *CTSZ* transcript by genotype of remaining SNPs can be found in **Figure S3D**-**F**. (**E**) Brightfield (BF) images and immunofluorescent staining of CTSZ and CD68 within a granuloma biopsy from an individual with pulmonary TB. Goat (Gt) and mouse (Ms) IgG isotype control staining is depicted in the top row. DAPI staining indicates cell nuclei. Images were captured at 100X magnification.

**Table 1:** *CTSZ* SNPs significantly associated with TB severity in Ugandan household contact study cohorts, sorted by ascending p-value. Included SNPs were significantly associated with Bandim TBScore after Bonferroni correction for multiple comparisons (p<0.05). Complete collection of 81 SNPs can be found in **Table S1**. SNPs are annotated as described in McHenry et al., 2023 [61]. Allele effects were assessed using a linear mixed effect model in the R package kimma to account for sex, HIV status, and genotypic principal component 1 and 2. Cohort 1 and 2 are independent cohorts of culture-confirmed adult TB cases. Abbreviations: SNP, single nucleotide polymorphism; CHR, chromosome; BP, base pair from GRCh38 build; Adj., adjusted; MAF, minor allele frequency.

| SNP | CHR:BP | Effect Allele | Adj. p-value | $\beta$ | MAF in Cohort 1 (n=149) | MAF in Cohort 2 (n=179) |
| --- | --- | --- | --- | --- | --- | --- |
| rs113592645 | 20:59001340 | T | 0.0001814 | -1.0036 | 0.18 | 0.061 |
| rs111630627 | 20:59002589 | G | 0.0003077 | -0.9268 | 0.18 | 0.075 |
| rs138964736 | 20:59002671 | ACTTTG | 0.0003077 | -0.9268 | 0.18 | 0.075 |
| rs76687632 | 20:59002905 | G | 0.0003077 | -0.9268 | 0.18 | 0.075 |
| rs8120779 | 20:59001977 | G | 0.0003942 | -0.8671 | 0.18 | 0.095 |

### CTSZ is produced in macrophages associated with human pulmonary granulomas

The human granuloma is an organized structure that can develop within the host to contain and restrict *Mtb* infection and is composed of heterogeneous immune cell populations, predominantly macrophages [64]. To investigate whether CTSZ production in macrophages is conserved between mice and humans, we performed immunostaining on granulomas biopsied from the lungs of patients with culture-confirmed TB. We positively identified CTSZ within TB granuloma-associated CD68^+^ macrophages from this lung tissue (**Figure 4E**). Thus, CTSZ is produced at the site of host-pathogen interaction in humans, suggesting that native functions at this interface could be interrupted should CTSZ production or localization be impeded. Combined with the results from *Ctsz* null mice, these data suggest that balancing cathepsin Z levels is required to regulate lung inflammation and reduce risk of mortality. Collectively, these data establish an association between human *CTSZ* variants and TB disease severity and reveal CTSZ as a granuloma-associated protein in human lungs for the first time.

## Discussion

Over fifteen years have passed since the initial discovery that human *CTSZ* is linked with TB disease susceptibility in West and South Africa. However, the relationship between *Mtb* susceptibility and CTSZ had yet to be experimentally determined. We show that genetic interruption of *Ctsz* in mice causes a failure of bacterial restriction and overproduction of CXCL1 during early *Mtb* infection, precipitating an increased risk of mortality. We further show a cell-autonomous effect of CXCL1 overproduction in macrophages during *Mtb* infection. We report 4 *CTSZ* SNPs and 1 INDEL significantly associated with TB severity in Ugandan individuals and show higher *CTSZ* expression in infected monocytes from this cohort. Finally, we find that CTSZ protein is present within the CD68^+^ macrophages in human granulomas, the pulmonary structure that contains and restricts *Mtb* growth.

CTSZ participates in several known immunological pathways [29,32,33,65]. Here, we show that lung CXCL1 levels are consistently elevated in *Ctsz^−/−^* mice prior to and throughout infection. Moreover, compared to wildtype siblings, *Ctsz^−/−^* macrophages produce more CXCL1 in response to pathogenic and non-pathogenic mycobacterial infection, suggesting a broad immunological response to bacterial exposure. CXCL1, a cytokine associated with severe TB disease in mice [51] and in humans [58], is primarily known as a neutrophil chemoattractant. Both *Ctsz* and *Cxcl1* are induced in *Mtb*-infected mice [66], but, to the best of our knowledge, this is the first study to directly link CTSZ and CXCL1 during TB pathogenesis. Future studies are needed to delineate the implications of this CTSZ-CXCL1 axis with other known roles of CTSZ, including cellular adhesion, migration, and antigen presentation [29]. For example, CTSZ is known to interact with cell surface integrins that mediate immune cell activity, including lymphocyte function-associated antigen-1 (LFA-1) [32,33] and macrophage-1 (Mac-1) antigen [65], which regulates *Mtb* phagocytosis [67] and phagocyte migration.

Deeper understanding of how the functions of CTSZ impact disease severity could prove vital to developing therapeutic strategies for both endogenous and infectious diseases beyond TB. For example, CTSZ has been implicated as a mediator of host response during *Helicobacter pylori* infections of patient-derived monocytes and *Salmonella* Typhimurium infections of murine BMDMs [68,69]. Further, mouse and human studies have investigated CTSZ for roles in aging [70,71] and in a number of endogenous conditions, including multiple sclerosis [72], primary biliary cholangitis [73,74], osteoporosis [75], and Alzheimer’s [76]. *CTSZ* has also been explored for prognostic value and roles in tumor progression across many cancers [77], including breast [78], colorectal [79], gastric [68], and prostate cancers [80], and hepatocellular carcinoma [81]. Increased *CTSZ* expression was associated with poor patient prognoses in some studies [79,81], with one study proposing *CTSZ* as a putative oncogene [81]. Given the importance of CTSZ in a spectrum of human disease categories, continued study of *CTSZ* may yield insights on the human response to departures from immune homeostasis.

While much remains unknown about the molecular roles of CTSZ during *Mtb* infection, this study is the first to identify cathepsin Z as a molecular correlate of TB severity in mice and humans. This study is also the first to report CTSZ localization within granuloma-associated CD68^+^ macrophages in *Mtb*-infected human lungs. Host genetic diversity is a central predictor of TB severity, and consideration of genetic diversity is essential to combat human pathogens as enduring and prolific as *Mtb*.

## Data Availability Statement

Relevant data is included in the manuscript and supplemental files. Additional data supporting this manuscript will be made available upon request.

## Acknowledgements

The authors acknowledge Martin Ferris, Rachel Lynch, and Ginger Shaw for coordination of experimental cohorts of CC mice through the UNC Systems Genetics Core Facility (SGCF); Christopher Sassetti, Douglas Marchuk, and Craig Lowe for thoughtful critiques and suggestions; and the staff of the Regional Biosafety Laboratory at Duke University for ongoing support of personnel safety in BSL-3 biocontainment. We acknowledge the staff of the Duke University BioRepository & Precision Pathology Center (BRPC) for identifying the human clinical cases and collecting and preparing the paraffin tissue sections. The authors also acknowledge the vital contribution of the patients whose samples provided data for this paper and the contributions made by senior physicians, medical officers, health visitors, laboratory personnel, and data personnel working with the Uganda-CWRU Research Collaboration. This study would not be possible without the generous participation of Ugandan patients and families.

## Funder Information

This work was funded by an NIH Director’s New Innovator Award AI183152 (C.M.Sm.), a Pew Scholars award (C.M.Sm.), and the following NIH grants: AI166304 (D.M.T.), AI127715 (D.M.T. and C.M.Sm.), AI181898 (T.R.H. and C.M.Sm.), AI162583 (T.R.H., C.M.St., P.H.B., H.M.-K.), N01-AI95383 (C.M.St.), and T32HL007567 (M.L.M.). M.T.P.G. was supported by a grant (88881.625374/2021-01) from the Fulbright Association and the Coordenação de Aperfeiçoamento de Pessoal de Nível Superior (CAPES). Research reported in this publication was supported in part by the Duke University Center for AIDS Research (CFAR), an NIH funded program (5P30 AI064518). The Duke University BRPC is supported in part by the NIH (P30CA014236). Biocontainment work performed in the Duke Regional Biocontainment Laboratory received partial support for construction and renovation from NIAID (UC6-AI058607 and G20-AI167200) and facility support from the NIH (UC7-AI180254). The sponsors or funders did not play any role in the study design, data collection and analysis, decision to publish, or preparation of the manuscript.

## Conflict of Interest

The authors declare no conflicts of interest.

## Materials & Methods

### Ethics Statement

All animal studies were conducted in accordance with the guidelines issued in the Guide for the Care and Use of Laboratory Animals of the National Institutes of Health and the Office of Laboratory Animal Welfare. Mouse studies were conducted at Duke University using protocols approved by the Duke Institutional Animal Care and Use Committee (IACUC) (Animal Welfare Assurance #A221-20-11 and #A204-23-10) in a manner designed to minimize pain and suffering in *Mtb*-infected animals. Any animal exhibiting signs of severe disease was immediately euthanized in accordance with IACUC-approved endpoints. Use of patient samples was approved by the Duke University Medical Center Institutional Review Board (IRB) under Protocol #00107795 and the University of Washington IRB under Protocol STUDY00001537. Patient sample processing at Duke University was carried out by Drs. Jadee Neff, Charlie Pyle, and Jason Stout. The human genetic data were obtained from the Kawempe Community Health Study in Uganda, which was approved by the National HIV/AIDS Research Committee of Makerere University (Protocol #014) and the University Hospitals Cleveland Medical Center IRB (Protocol #10-01-25). Final clearance was given by the Uganda National Council for Science and Technology (Ref #658).

### Mice

Male and female C57BL/6J (#000664) and male B6.129S7-*Ifngr1^tm1Agt^>*/J (*Ifngr^−/−^*; #003288) mice were purchased from The Jackson Laboratory. Male CC033/GeniUncJ (CC033) and CC038/GeniUnc (CC038) mice were purchased from the University of North Carolina (UNC) Systems Genetics Core Facility (SGCF). *Ctsz^−/−^* mice were generously provided by Robin Yates (University of Calgary, Calgary, AB, Canada) [82]. *Ctsz^+/+^>* and *Ctsz^−/−^*mice were subsequently bred at Duke. All mice were housed in a specific pathogen-free facility within standardized living conditions (12-hour light/dark, food and water *ad libitum*). Aerosol-infected mice were matched at 8-12 weeks of age at the time of *Mtb* infection. Mice were individually identified for weighing and wellness assessment throughout infection using Bio Medic Data Systems implantable electronic ID transponders (TP-1000) implanted subcutaneously at the back of the neck prior to infection.

### Genotyping

In-house confirmation of *Ctsz^−/−^* genotype was performed using forward primer 5’-TTG CTG TTG GCG AGT GCG-3’ and reverse primer 5’-CTT GTC ACC AGA TTC CAG C-3’ to detect wildtype *Ctsz* and forward primer 5’-GCT ACC TGC CCA TTC GAC-3’ and reverse primer 5’-ACA GTA GGA CTG GCC AGC-3’ to detect knockout product. Primer sequences were generously provided by Robin Yates (University of Calgary, Calgary, AB, Canada). DNA was extracted from tissue samples using the DNEasy Blood & Tissue Kit (Qiagen). DNA products were prepared for PCR using Q5 High-Fidelity Master Mix (New England BioLabs) and amplified. Protocol included initial 98°C (30s), then 34 cycles of denaturation (98°C, 10s), annealing (68°C, 30s), and extension (72°C, 90s), and a final 72°C (180s), resting at 10°C ∞ until stopped. Amplified products were separated on a 1% agarose-TAE gel using SYBR Safe stain (Thermo Fisher Scientific) and 1kb Plus DNA Ladder (New England BioLabs). *Ctsz^+/+^>*and *Ctsz^−/−^* mice were genotyped at the time of weaning from ear and tail tissue biopsies by TransnetYX (Cordova, TN, USA) using proprietary RT-PCR primers designed to detect both *lacZ*, present in the IRES vector disturbing the second exon of *Ctsz* [82], and wildtype *Ctsz*.

### Bacterial Strains & Culture

All infections were performed with either *Mtb* H37Rv genotype or *M. bovis* BCG (Bacillus Calmette-Guérin) Danish (gift from Sunhee Lee, University of Texas Medical Branch, Galveston, TX, USA), which was transformed with pTEC-15 wasabi fluor and possesses a hygromycin resistance marker for selection [83]. Aerosol infections were performed using an *Mtb* H37Rv strain confirmed to be positive for the cell wall lipid and virulence factor phthiocerol dimycocerosate (PDIM; gift from Kyu Y. Rhee, Weill Cornell Medicine, New York, NY, USA). Bacteria were cultured in Middlebrook 7H9 medium supplemented with oleic acid-albumin-dextrose catalase (OADC), 0.2% glycerol, and 0.05% Tween 80 (or 0.005% tyloxapol for macrophage infections) to log-phase with shaking (200rpm) at 37°C. Hygromycin (50 µg/mL) was added when necessary. Prior to all *in vivo* infections, cultures were washed and resuspended in phosphate-buffered saline (PBS) containing 0.05% Tween 80 (hereafter PBS-T). Bacterial aggregates were then broken into single cells using a blunt needle before diluting to desired concentration for infection.

### Mouse Infections

Mice were infected with ∼150-350 *Mtb* CFU via aerosol inhalation (Glas-Col). On the day following each infection, one cage was sacrificed to enumerate lung CFU as an approximation of infectious dose. For all infections, mice were euthanized in accordance with approved IACUC protocols, and lung and spleen were harvested into PBS-T and processed in a FastPrep-24 Homogenizer (MP Biomedicals, 4.0m/s, 45s, 2-3x). *Mtb* burden was quantified by dilution plating onto Middlebrook 7H10 agar supplemented with OADC, 0.2% glycerol, 50 mg/mL Carbenicillin, 10 mg/mL Amphotericin B, 25 mg/mL Polymyxin B, and 20 mg/mL Trimethoprim. Lung homogenate was centrifuged through a 0.2µm filter to collect decontaminated filtrate, and cytokines and chemokines were assayed using the pro-inflammatory focused 32-plex assay (Eve Technologies, Calgary, AB, Canada).

### Human Tissue Immunofluorescent Staining

Patient tissue samples containing *Mtb* granulomas were identified at the Duke University School of Medicine. Clinical tissue specimens were obtained from the Duke Pathology Department, and 5µm paraffin sections for antibody staining were cut by the Research Histology Laboratory within the BioRepository & Precision Pathology Center (BRPC). Paraffin was dissolved using two xylene washes followed by washes with ethanol of increasing dilution (100% twice, 95% twice, 70% once, 50% once), three washes with deionized water, and a final wash in PBS. Sample was placed in antigen retrieval buffer (10 mM Tris/1 mM EDTA, pH 9.0) and processed in a pressure cooker for 10min. Following a cooling step, samples were blocked for an hour in 2.5% normal donkey serum. Samples were incubated overnight at 4°C with Goat anti-Human/Mouse/Rat Cathepsin X/Z/P Polyclonal Antibody (R&D Systems, AF934, 0.185 mg/mL) and Mouse anti-Human CD68 Monoclonal Antibody (Agilent Dako, M081401-2, 0.185 mg/mL) in 2.5% serum in a humidified chamber. Immunoglobulin G (IgG) isotype controls for background staining (Goat: Biotechne, AB-108-C, 1 mg/mL stock; Mouse: GenScript, A01007, 1 mg/mL stock; Rabbit: Invitrogen, 10500C, 3 mg/mL provided) were also used. Primary antibody was removed with three washes of PBS and two of deionized water. Samples were incubated in Alexa Fluor (AF) conjugated secondary antibody (Thermo Fisher Scientific, 1:500; Donkey anti-Goat IgG AF Plus 647: A32849; Donkey anti-Mouse IgG AF 488: A-21202; Donkey anti-Rabbit IgG AF 555: A-31572) in 2.5% serum for 1-3 hours. Following three PBS washes, the samples were mounted for imaging in DAPI Fluoromount-G (Southern Biotech, 0100-20) on glass slides (Fisher Scientific, 22-035813). All antibodies used for staining were centrifuged at 10000 RCF (4°C) for 10min to remove antibody precipitate prior to use.

### Microscopy Analysis

Human samples were imaged at 100X on a Zeiss Axio Observer Z1 inverted microscope with an X-Light V2 spinning disk confocal imaging system (Biovision). Images were processed identically within Fiji software (v2.14.0/1.54f) for image clarity.

### Bone Marrow-Derived Macrophage Infections

*Ctsz^+/+^>* and *Ctsz^−/−^* sibling pairs were sacrificed in accordance with approved IACUC protocols between 10-12 weeks of age. For BCG infections, bone marrow was flushed from hip and leg bones with DMEM (Corning) and cultured for a week at 37°C in a sterile solution of DMEM with 10% heat-inactivated fetal bovine serum (Corning), 18% 3T3-derived M-CSF, 1X Pen/Strep (Corning), and 25mM HEPES (gibco). For *Mtb* infections, bone marrow was flushed from hip and leg bones with sterile DMEM (Corning) and frozen in 10% DMSO in heat-inactivated fetal bovine serum (Corning). Aliquots were later thawed and cultured for a week at 37°C in a sterile solution of DMEM with 10% heat-inactivated fetal bovine serum (Corning), 30 µg/mL recombinant M-CSF (PeproTech), 1X Pen/Strep (Corning), and 25mM HEPES (gibco). Differentiated macrophages were then plated at a concentration of 4×10^5^ cells/well in a 24-well plate and cultured at 37°C overnight in a DMEM solution as above but without Pen/Strep. BMDMs were infected with BCG or transported to BSL-3 biocontainment for infection with *Mtb* at MOI 3 or 7 or left uninfected. Wells were tested for even infection by CFU plating. At 24 hours post-infection, supernatants were collected and filtered using a 0.2µm filter to remove bacteria. Cytokines and chemokines were assayed from using the high-sensitivity 18-plex discovery assay (Eve Technologies, Calgary, AB, Canada).

### Western Blotting

To compare the presence of mouse CTSZ in *Ctsz^+/+^>* and *Ctsz^−/−^* mice, protein was extracted from lung and spleen homogenate (12-14 weeks of age) using RIPA buffer (Sigma-Aldrich) and 1X Protease Inhibitor Cocktail (Sigma-Aldrich) after a PBS wash. For the comparison of mouse CTSZ between uninfected B6 mice and *Tip5^S^>* CC strains (CC033 and CC038), whole lungs were collected from male mice (8 weeks of age) into 1mL of Trizol reagent. Samples were homogenized with sterile beads at 4.5 m/s for 30s using the FastPrep-24 Homogenizer (MP Biomedicals). For samples in Trizol, protein was precipitated for 15min using 9 volumes of 100% methanol at room temperature. The protein precipitate was centrifuged at 3000rpm for 5min, dried for 5min, and washed in an equal volume of 90% methanol. The protein precipitates were then centrifuged for 1min at 3000rpm, dried for 10min, resuspended in 1mL of RIPA buffer and 1X Protease Inhibitor Cocktail, and heated for 5-10min at 95°C. Equal volumes of each sample were combined with Laemmli Sample Buffer (BioRad) and 2-Mercaptoethanol (BioRad) and heated at 95°C for 5min. SDS-PAGE was performed using BioRad Western Blotting kit along with Precision Plus Protein All Blue Prestained Protein Standards (BioRad). Protein was separated using a 4-20% Mini-PROTEAN TGX Stain-Free Protein Gel (BioRad) and transferred to a polyvinylidene fluoride (PVDF) membrane using a semi-dry transfer protocol on a Trans-Blot Turbo Transfer System (BioRad). Membrane was blocked using EveryBlot Blocking Buffer (BioRad). Primary staining was performed at 4°C overnight using Human/Mouse/Rat Cathepsin X/Z/P Antibody (R&D Systems; AF934; 1:2000 dilution in EveryBlot Blocking Buffer). For B6 and *Tip5^S^>* CC mice, 0.1% Tween 20 was added to the blocking buffer and primary staining also included Vinculin (E1E9V) XP Rabbit mAb (Cell Signaling; #13901; 1:5000 dilution in EveryBlot Blocking Buffer + 0.1% Tween 20). For *Ctsz^+/+^>* and *Ctsz^−/−^* mice, secondary staining was performed at room temperature for 60min using Donkey anti-Goat 680 (LI-COR; 1:20000 dilution in EveryBlot Blocking Buffer + 0.1% SDS). Blot was washed in TBS-T between blocking and antibody stains, and fluorescence was measured using a LI-COR Odyssey. Secondary staining for B6 and *Tip5^S^>* CC mice was performed at room temperature for 60min using HRP-conjugated Rabbit Anti-Goat IgG (Proteintech; SA00001-4; 1:5000 dilution in EveryBlot Blocking Buffer + 0.1% Tween 20) and HRP-conjugated Goat Anti-Rabbit IgG (Proteintech; SA00001-2; 1:5000 dilution in EveryBlot Blocking Buffer + 0.1% Tween 20). Blot was washed in PBS-T (0.1% Tween 20). Chemiluminescence was developed using SuperSignal West Pico PLUS Chemiluminescent Substrate (Thermo Fisher Scientific) and imaged using a ChemiDoc Plus Imaging System (BioRad). Quantification of the blot was performed with ImageLab software (version 6.1).

### Human CTSZ Analysis

We queried the summary statistics from a published genome-wide association study (GWAS) of TB severity in cases from Kampala, Uganda [61]. Briefly, two independent cohorts of culture-confirmed adult TB cases (n=149, n=179) [84] were included in a GWAS. TB severity was quantified using the Bandim TBscore, which enumerates TB symptoms (e.g., cough, hemoptysis, dyspnea) and clinical signs (e.g., anemia, low body mass index, high body temperature) [62,85]. SNPs within *CTSZ* were identified using a 5kb flanking region around the *CTSZ* start and end positions reported in Ensembl (GRCh38). Pairwise linkage disequilibrium (LD) for these SNPs was evaluated as the squared inter-variant allele count correlations (R^2^>) using PLINK (version 1.90) in the larger of the two cohorts (n=179). An LD plot was generated from these pairwise LD measures using the R package LDheatmap (version 1.0-5) [86]. The model used to estimate allele effects accounted for sex, HIV status, RNA-Seq batch, and genotypic principal components 1 and 2.

SNP eQTL assessment was performed for the four significant SNPs indicated in **Table 1**. A linear mixed effect model was developed in the R package kimma [87] to compare baseline, media, and *Mtb*-induced *CTSZ* expression against each SNP genotype. The eQTL model accounted for sex, HIV status, RNA-Seq batch, and genotypic principal components 1 and 2. *CTSZ* expression as log_2_(counts per million) was obtained from RNA-Seq data normalized using voom [88]. RNA-Seq data used for these analyses originated from a previously published dataset of CD14^+^ monocytes isolated from individuals within the Uganda cohort [63]. Monocytes were subjected to 6-hour media or *Mtb* stimulation and transcriptionally assayed.

### Statistical Analysis & Data Visualization

Hypothesis testing was performed using R statistical software (version 4.3.1). Statistical tests used for hypothesis testing are noted in figure legends. Shapiro-Wilks tests were used to assess normality in phenotype data prior to parametric hypothesis testing, and log_10_-transformation was applied for normalization where appropriate. Kaplan-Meier survival curves were calculated using the R package survminer (version 0.5.0). A visualization of mouse chromosome 2 was generated using the R package karyoploteR (version 1.16.0) from the GRCm38/mm10 mouse genome build. Heatmaps in **Figure 1A** and **Figure 2C** were generated using the R packages ComplexHeatmap (version 2.21.2) and heatmaply (version 1.5.0), respectively. Optimization and sparse partial least squares discriminant analysis (sPLS-DA) on time course infection cohorts were performed on timepoint data using the R package mixOmics (version 6.24.0).

## Supplemental Table Legend

**Table S1: Complete list of 81 *CTSZ* SNPs present in Ugandan household contact study cohorts and their associations with TB severity**. TB severity was evaluated by Bandim TBScore. Summary statistics for the *CTSZ* variants shown are based on a meta-analysis of two independent cohorts of culture-confirmed adult TB cases (described in McHenry et al., 2023 [61]). Each cohort utilized a linear regression model that controlled for HIV status, sex, and one principal component. Unadjusted p-values are reported. Abbreviations: CHR, chromosome; BP, base pair from GRCh38 build; MAF, minor allele frequency.

## Notes

### Competing Interest Statement

The authors have declared no competing interest.

