## Supplemental Figures 1-3 for "Cathepsin Z is a conserved susceptibility factor underlying tuberculosis severity"

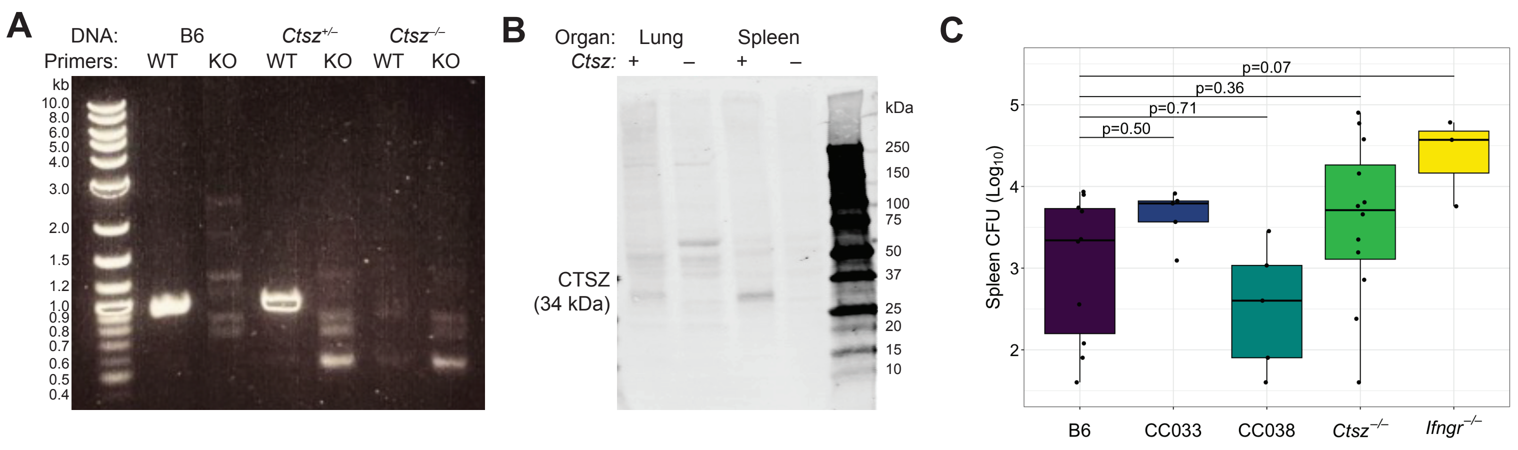


**Figure S1: Genetic validation and infection of *Ctsz*$^{-/-}$ mice.** (**A**) Expression of wildtype and truncated *Ctsz* in tail sections from B6, *Ctsz*$^{+/-}$, and *Ctsz*$^{-/-}$ mice. (**B**) CTSZ protein in lung and spleen homogenate of *Ctsz*$^{+/+}$ and *Ctsz*$^{-/-}$ mice. (**C**) Bacterial burden measured by dilution plating from spleen homogenate 4 weeks after aerosol infection with *Mtb* H37Rv (n=3-12 per strain; all males except B6 and *Ctsz*$^{-/-}$ groups, which included both sexes in equal proportion). Hypothesis testing was performed by one-way ANOVA and Dunnett's *post hoc* test on $\log_{10}$-transformed values.

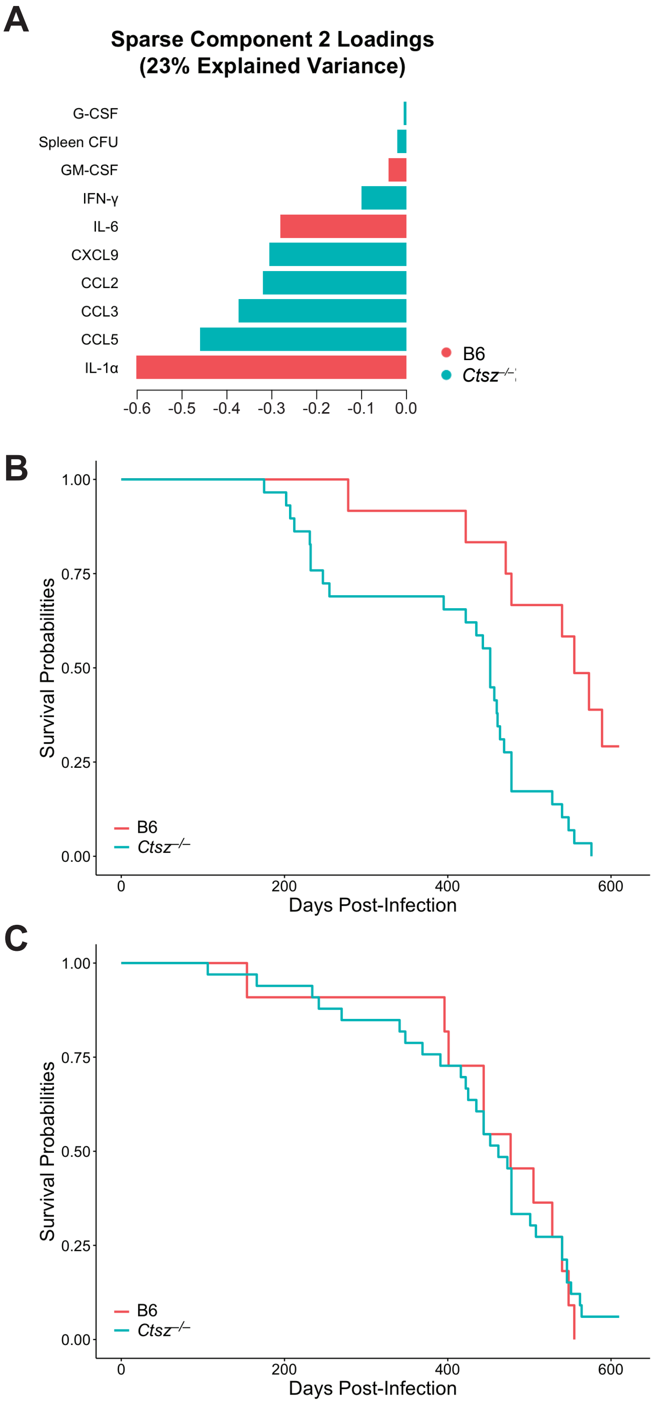


**Figure S2: sPLS-DA and survivial analysis comparing *Ctsz*$^{-/-}$ and B6 mice.** (**A**) Phenotype loadings contributing to sparse component 2. Mice were sacrificed at 2, 3, 4, and 8 weeks after aerosolized *Mtb* infection. Data are from two experiments with n=6-14 mice per genotype, representative of both sexes, at each timepoint. Kaplan-Meier survival estimates of aerosol-infected B6 (n=23) and *Ctsz*$^{-/-}$ mice (n=62) across two independent experiments, among (**B**) male (p=3e-04) and (**C**) female (p=0.9) mice. Equal proportions of both sexes included.

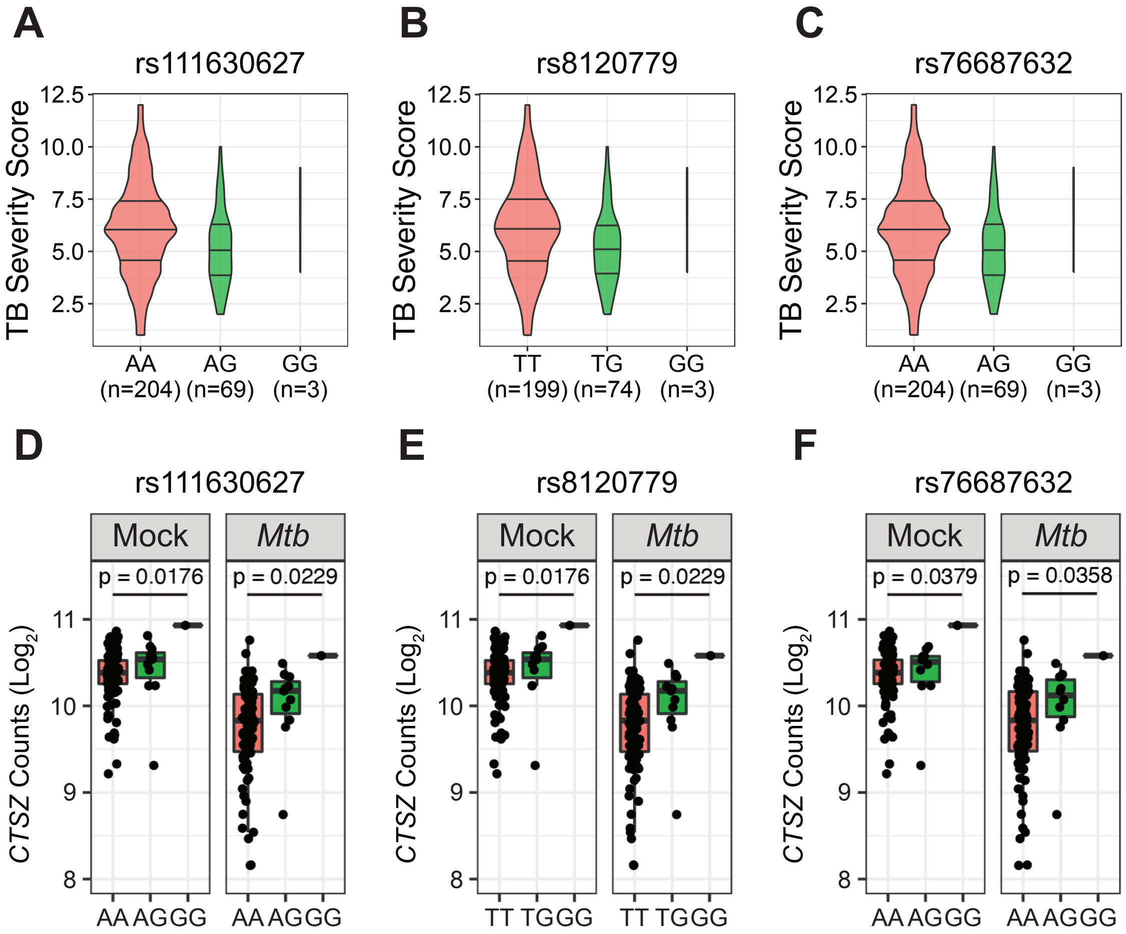


**Figure S3: Minor alleles of *CTSZ* SNPs within the TB severity haplotype block are associated with lower TB severity score and significantly greater *CTSZ* expression**. Comparison of TB severity, measured using Bandim TBScore, by genotype for (**A**) rs111630627, (**B**) rs8120779, and (**C**) rs76687632 SNPs. Expression of each allele of each SNP was assessed by RNA-Seq at 6 hours after mock and *Mtb* infection in human-derived monocytes. *CTSZ* expression by monocytes harboring the minor allele for each SNP was significantly increased following both infection conditions for the (**D**) rs111630627, (**E**) rs8120779, and (**F**) rs76687632 SNPs. eQTL effects were assessed with a linear mixed effect model in kimma to account for sex, age, RNA-Seq batch, genotypic principal components 1 and 2, and kinship.
